# movedesign: Shiny R app to evaluate sampling design for animal movement studies

**DOI:** 10.1101/2023.01.27.525894

**Authors:** Inês Silva, Christen H. Fleming, Michael J. Noonan, William F. Fagan, Justin M. Calabrese

## Abstract

1. Projects focused on movement behavior and home range are commonplace, but beyond a focus on choosing appropriate research questions, there are no clear guidelines for such studies. Without these guidelines, designing an animal tracking study to produce reliable estimates of space-use and movement properties (necessary to answer basic movement ecology questions), is often done in an *ad hoc* manner.
2. We developed ‘`movedesign`’, a user-friendly Shiny application, which can be utilized to investigate the precision of three estimates regularly reported in movement and spatial ecology studies: home range area, speed, and distance traveled. Conceptually similar to statistical power analysis, this application enables users to assess the degree of estimate precision that may be achieved with a given sampling design; *i*.*e*., the choices regarding data resolution (sampling interval) and battery life (sampling duration).
3. Leveraging the ‘`ctmm`’ `R` package, we utilize two methods proven to handle many common biases in animal movement datasets: autocorrelated Kernel Density Estimators (AKDE) and continuous-time speed and distance (CTSD) estimators. Longer sampling durations are required to reliably estimate home range areas via the detection of a sufficient number of home range crossings. In contrast, speed and distance estimation requires a sampling interval short enough to ensure that a statistically significant signature of the animal’s velocity remains in the data.
4. This application addresses key challenges faced by researchers when designing tracking studies, including the trade-off between long battery life and high resolution of GPS locations collected by the devices, which may result in a compromise between reliably estimating home range or speed and distance. ‘`movedesign`’ has broad applications for researchers and decision-makers, supporting them to focus efforts and resources in achieving the optimal sampling design strategy for their research questions, prioritizing the correct deployment decisions for insightful and reliable outputs, while understanding the trade-off associated with these choices.

## 1. Background

Modern tracking technologies are advancing the fields of ecology and other fields of biology, with data-rich studies revealing how movement shapes ecological processes, biological interactions, and behavioral responses to natural or human-induced environmental changes (Kays et al., 2015; Nathan et al., 2022). Biologging and telemetry devices, such as global positioning system (GPS) and very-high frequency (VHF) transmitters, are common sampling methods in this field, and have been used to answer a multitude of research questions ―the majority of which rely on estimates of large-scale processes, such as home ranges (Fleming & Calabrese, 2017; Horne et al., 2019), or fine-scale processes, such as speed or distance traveled (Gurarie et al., 2016; Noonan, Fleming, et al., 2019).

The technology is improving, with cheaper, less invasive tags available (Foley & Sillero-Zubiri, 2020; Gottwald et al., 2019), allowing researchers access to high-volume high-resolution movement data for an increasing number of species (Kays et al., 2015): the animal tracking data platform Movebank (www.Movebank.org) already hosts 2.4 billion locations for over 1,000 species. Although these data are critical for conservation, tracking devices can be costly and/or logistically difficult to deploy (Hays et al., 2016), requiring us to weigh their benefits against their costs to ensure optimal outcomes. VHF transmitters —here referring to their use in triangulation or homing of tracked animals (White & Garrott, 2012)— remain a cost-effective method for data collection on small animals, as well as most reptiles, due to constraints in weight and/or tag attachment (such as surgical implantation).

Researchers often track as many animals as possible, for as long as possible, recording new locations at regular, short sampling intervals —particularly if the goal is to identify finer movement behaviors (Nathan et al., 2022). However, although the use of GPS loggers typically allows for locations to be recorded at higher rates than VHF transmitters (which, without access to automated methods, would require triangulation/homing to be performed regularly every few seconds/minutes for high-frequency data), achieving both long battery life and high sampling frequency is challenging. Excluding other factors that may further reduce the longevity of tracking devices in the wild (*e*.*g*., additional sensors, signal acquisition time, algorithm efficiency, data recovery method), GPS battery life shortens with shorter sampling intervals (Forrest et al., 2022; Moriarty & Epps, 2015). Successfully designing an animal tracking study therefore requires a compromise between *sampling duration* (how long an animal should be tracked for) and *sampling interval* (time between which new locations are recorded; reciprocal of sampling frequency). Moreover, target analyses and sampling design are tightly linked, so we must keep specific goals in mind: there are circumstances where we may be able to address either large-scale or fine-scale questions, but not both concurrently.

Our objective is to provide a more rigorous *a priori* approach, analogous to statistical power analysis (Steidl et al., 1997), for determining the ideal sampling duration and sampling interval of an animal tracking project. We consider three common estimates— home range area, speed, and distance traveled—through the lens of continuous-time movement models and the ‘`ctmm`’ `R` package (Calabrese et al., 2016). The ‘`ctmm`’ methods are insensitive to the sampling schedule and deal with many of the biases present in modern movement datasets (such as those due to autocorrelation, small sample sizes, irregular or missing data, or measurement error), facilitating robust comparisons across individuals, species, and ecosystems (Fleming et al., 2015, 2018, 2019; Fleming & Calabrese, 2017; Silva et al., 2022). However, like many statistical tools, these methods still require adequate sample sizes to achieve high accuracy in their outputs; for example, pHREML-AKDE estimates (designed to handle small effective sample sizes) still tend to be negatively biased with effective sample sizes below 30 (Silva et al., 2022). Furthermore, like all proper estimators, statistical uncertainty decreases with increasing (effective) sample sizes. We chose to implement this in the Shiny `R` application, ‘`movedesign`’ (current version 0.1.3) to facilitate adoption and accessibility to the broadest possible audience, and hope that we can contribute to the increase of comparable and reliable movement ecology studies. We note that, for most use cases, this application may require existing datasets. This application was built using the ‘`golem`’ framework (Fay et al., 2021), and is available as an installable `R` package at https://github.com/CASUS/movedesign.

## 2. Conceptualization

### 2.1. Parameters

Explicitly modelling all the complex interactions between internal states, biophysical constraints, environmental cues and social dynamics that determine animal movement is generally considered impossible (Codling et al., 2008). However, by interpreting movement as a continuous-time stochastic process (Calabrese et al., 2016; Fleming et al., 2014a; Gurarie & Ovaskainen, 2011), we can efficiently summarize behavior using characteristic timescales, including the position autocorrelation (*τ*_*p*_) and velocity autocorrelation (*τ*_*v*_) timescale parameters, and describe long-term dispersal behavior and short-term movement behavior across levels of data resolution and duration. The autocorrelation timescales indicate the time required for the correlations in the focal quantity (here position or velocity) to decay by a factor of 1/*e* —conventionally, the data is considered effectively independent when 5% or less autocorrelation remains (Gurarie & Ovaskainen, 2011). These timescales impose constraints on the sampling design that must be met in order to evaluate both small-scale (speed and distance) and large-scale phenomena (home range). (Fleming et al., 2014b). If these constrains are not met, it could lead to biases and estimate uncertainty.

The position autocorrelation parameter (*τ*_*p*_) can be interpreted as the home range crossing time, or the time is takes on average for an animal to cross the linear extent of its range (Silva et al., 2022). As *τ*_*p*_ increases, we can expect an animal to take longer to travel this linear extent. Range-resident animals tend to travel away from their point of origin at a rate controlled by the location variance parameter (*σ*_*p*_) while simultaneously reverting back to it at a rate driven by *τ*_*p*_ (Péron et al., 2017). As *τ*_*p*_ → ∞, movement becomes endlessly diffusive (*σ*_*p*_ → ∞), with no range residency behavior; for range-resident species, *σ*_*p*_ is asymptotically constant and proportional to home range area (Calabrese et al., 2016; Fleming et al., 2014a).

The velocity autocorrelation parameter (*τ*_*v*_) describes how velocity persists through time (directional persistence; Fleming *et al*. 2014). Animals with strong directional persistence (longer bouts of constant speed and constant direction) will tend to have a large *τ*_*v*_ parameter (such as a migratory species), while animals with more tortuous movement (less linear, more diffusive) tend towards smaller *τ*_*v*_. Speed and distance traveled are two of the properties of an animal’s velocity process, with variance *σ*_*v*_ = *σ*_*p*_/(*τ*_*p*_ × *τ*_*v*_) for stationary processes (Noonan, Fleming, et al., 2019). For any value of *σ*_*v*_, as *τ*_*v*_ → ∞, the movement process approximates ballistic motion.

### 2.2. Sampling design

Study design optimization requires clearly defined research questions and objectives (Fieberg & Börger, 2012), and an understanding of existing constraints, both those related to the study species (characterized by *τ*_*p*_, *τ*_*v*_, *σ*_*p*_, and *σ*_*v*_) and the sampling parameters (duration, and interval).

Our choice of sampling parameters largely determines our ability to detect the characteristic timescales of the movement process: it is the sampling duration and interval relative to characteristic timescales that determine whether there will be any signature of the animal’s range crossing time (*τ*_*p*_), or its directional persistence (*τ*_*v*_). Typically, sampling duration (*T*) should be at least as long as *τ*_*p*_ (and ideally many times longer) for home range estimation (Fleming et al., 2019; Fleming & Calabrese, 2017; Silva et al., 2022). For estimating speed and distance, the sampling interval (Δ*t*) needs to be less than or equal to the velocity autocorrelation timescale for distance and speed estimation, *i*.*e*., Δ*t* ≤ *τ*_*v*_ (Noonan, Fleming, et al., 2019). If Δ*t* > 3*τ*_*v*_ then no statistically significant signature of the animal’s velocity will remain (Noonan, Fleming, et al., 2019). We can evaluate how much information is present in an animal tracking dataset by considering the *effective sample size* (*N*). This concept, while well-discussed in statistical literature, is frequently ignored in movement ecology studies. *Effective sample sizes* are the number of independent locations that provide the same information as the non-independent data under consideration (Fieberg & Börger, 2012; Fleming et al., 2019; Silva et al., 2022) —and differ depending on the parameter under consideration. For the purposes of home range estimation, if an animal crosses its home range once per day and we collect locations for 100 days, our effective sample size (*N*) will be roughly 100. This equivalence is irrespective of sampling interval Δ*t* (provided that Δ*t* < 1 day), as *N* here is driven by the number of days sampled, and not by the number of samples per day. Simply put, the information present in this autocorrelated tracking dataset will not change if we collected new locations once a day (for an absolute sample size, *n*, of 100 locations), every hour (*n* = 2,400 locations), or every minute (*n* = 144,000 locations). Considering that recorded locations are frequently autocorrelated in modern tracking datasets (and that most species take longer than a day to cross their home range; Calabrese *et al*. 2016; Patterson *et al*. 2017), obtaining a satisfactory effective sample size (*N* > 30) can be challenging but is essential for a reliable movement metric or space-use estimate. For example, if the study species takes months to cross its home range, as is the case of the Mongolian Gazelle (average τ_p_ of 6.6 months; Fleming et al., 2014b), achieving an *N* = 30 could require an individual to be tracked for over 16 years, which is substantially longer than their average lifespan.

The choice between tracking devices further complicates sampling design. For GPS/satellite loggers, as sampling is automated and conducted by satellite systems, battery life is inherently linked to both sampling duration and frequency —*i*.*e*., choosing a higher fix rate (shorter intervals between recorded locations) leads to lower battery life (and, therefore, shorter sampling durations); excluding self-charging models such as solar tags, researchers are required to choose one over the other. For Very-High-Frequency (VHF) transmitters, battery life is decoupled from any specific sampling interval, and depends only on the sampling duration specified by the manufacturer —*i*.*e*., researchers choose a transmitter based on its battery life, but the interval between new locations is subject only to the data collection employed manually in the field (locating the animal every hour, or every day, has no influence on the battery life)..

In contrast with conventional radio-telemetry, GPS loggers have a higher potential to resolve fine-scale movements; however, the trade-off between battery life and resolution is more conspicuous, and with a greater impact on potential outputs. To facilitate sampling design within ‘`movedesign`’, we visualize this trade-off (sampling duration *versus* sampling interval) by simulating GPS battery life as a three-parameter log-logistic function (Appendix S1), based on the battery life-resolution trade-off of several GPS models of animal tracking devices.

### 2.3. Analytic scope

Our evaluation of sampling design focuses on the precision of three estimates at two different scales ―large-scale (*home range area*) and fine-scale (*speed* and *distance* traveled). As the choice of statistical estimator can further obscure estimate interpretation and comparison (Fieberg & Börger, 2012; Fleming et al., 2015), we leverage the ‘`ctmm`’ package to develop a reliable inferential framework for sampling design evaluation, generating comparable estimates with confidence intervals (Calabrese et al., 2016; Fleming et al., 2018, 2019; Fleming & Calabrese, 2017).

*Home range area* is one of the core outputs of movement or spatial ecology studies and provides information on the long term space-use requirements, which are invaluable for conservation policies and protected area delimitation (Bartoń et al., 2019; Lambertucci et al., 2014). Here, we follow the definition of home range as the area repeatedly used throughout an animal’s lifetime for all its normal behaviors and activities, excluding the occasional exploratory excursions (Burt, 1943; Calabrese et al., 2016; Silva et al., 2022), and estimate it using the Autocorrelated Kernel Density Estimator (AKDE; Calabrese *et al*. 2016), which accounts for the autocorrelation present in tracking data.

Both speed and distance traveled provide quantifiable links between behavior and energetics, as demographic outcomes and population dynamics are affected by how far and how fast animals must travel to meet their nutritional and reproductive needs (Morales et al., 2010; Noonan, Fleming, et al., 2019). *Distance* traveled is usually equated with the sum of the straight-line displacement (SLD) between all subsequent locations, while *speed* is the sum of SLD divided by time. Here, we use the continuous-time speed and distance (CTSD) estimator instead, as it overcomes important limitations of SLD estimation (Noonan, Fleming, et al., 2019). SLD tends to overestimate speed and distance traveled at small sampling intervals and underestimates these quantities at large sampling intervals (see Noonan, Fleming, et al., 2019).

There are two objective metrics that quantify the performance of these estimators: the effective sample size for home range estimation (*N*_*area*_), or how many independent location fixes would be required to calculate the same quality home range estimate, and the effective sample size for speed estimation (*N*_*speed*_), or how many independent velocity fixes would be required to calculate the same quality mean speed estimate.

In addition, the accuracy and precision of the estimates above —calculated from any sampling design— is judged by the relative error of both point estimates compared to the exact expectation values from simulations, and their confidence intervals. For home range areas, we derive the exact expectation value of the true 95% area based on the model from which data are simulated (see Fleming et al., 2014b; Noonan, Tucker, et al., 2019; Silva et al., 2022). For speed and distance traveled, the expected values are extracted from a fine-scale error-free trajectory simulated for 10 days at a sampling interval of *τ*_*v*_/10 (Noonan, Fleming, et al., 2019), functioning as a Riemann sum approximation of the path length that achieves a relative error of 𝒪(10^−3^) while reducing computation time.

## 3. Workflow

The main goal of ‘`movedesign`’ is to determine if a particular sampling design is sufficient to answer an *a priori* defined research question, following the conceptual workflow presented in Figure 1. First, users should consider a proxy dataset (typically the same species, or one with similar movement behaviors; see below for further details) from which to extract the aforementioned measures: characteristic timescales (τ), location variance (*σ*_*p*_), and velocity variance (*σ*_*v*_). These measures are extracted during automated model fitting and selection (Fleming et al., 2019) —a step that allows users to proceed with study design experimentation without requiring prior knowledge of the underlying movement process. If known *a priori*, users can also directly input these measures and skip the extraction step. Second, we input sampling design parameters ― duration (*T*) and interval (Δ*t*) ― to generate simulated datasets conditional on the previously fitted model. Third, we run estimators for home range and/or speed/distance traveled on the simulated data, while calculating the expected values in the background, from the model and parameters used in the simulation. Finally, we compare the estimates with the expected values (“truth”) and with simulation results from previous studies (Noonan, Fleming, et al., 2019; Silva et al., 2022), providing an overview of the reliability of that sampling design for estimating home range areas or speed and distance traveled.

**Figure 1.**
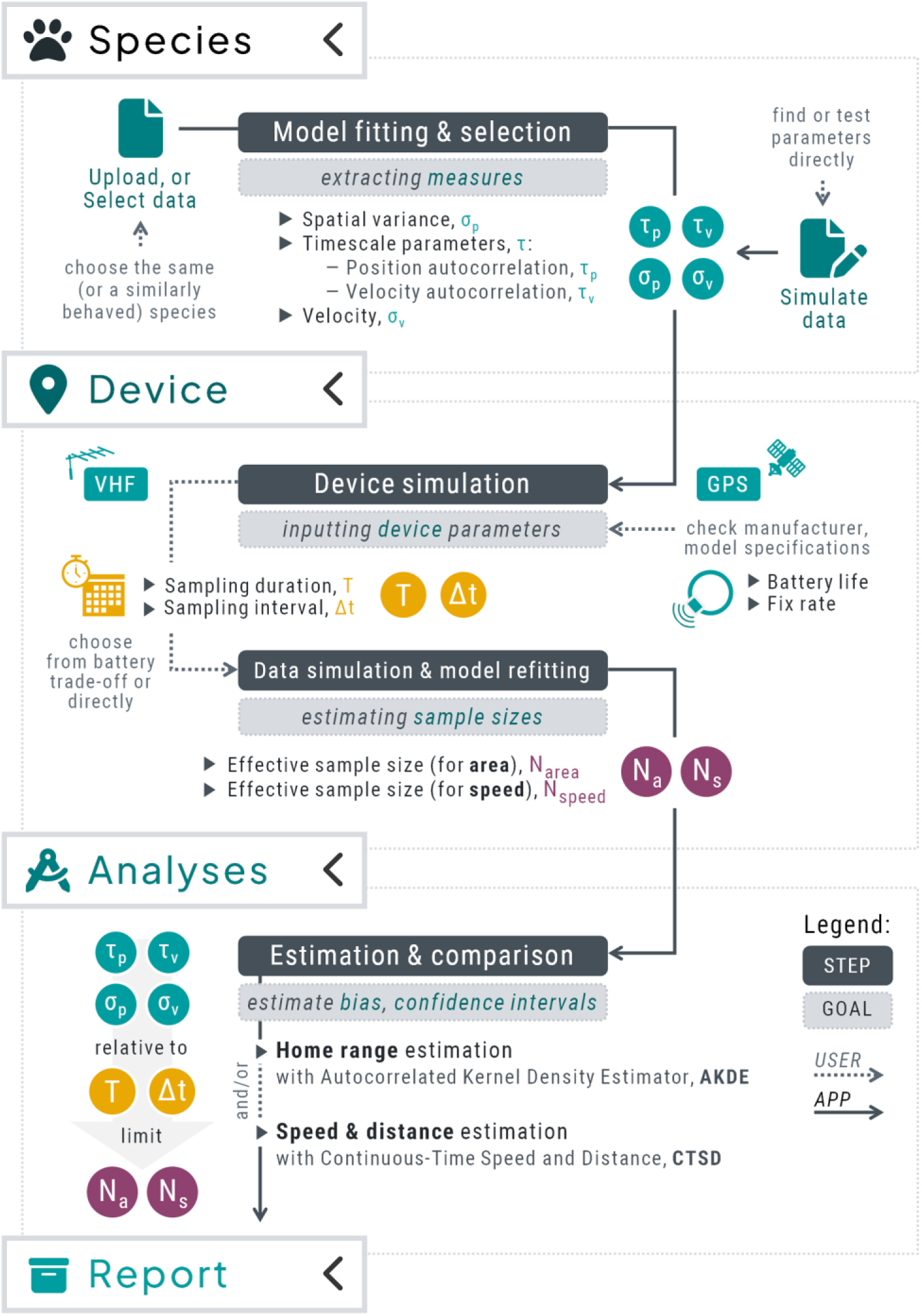
Conceptual workflow of the key elements of the `‘movedesign’` application, providing a R/Shiny-powered user interface to test different sampling designs for specified tracking projects. The application is divided into four main sections: “Species” (where data are uploaded, selected or simulated to extract relevant parameters and measures), “Device” (where users input device parameters, visualize potential trade-offs, and evaluate effective sample sizes), “Analyses” (where users can estimate home range areas, and speed and distance traveled,, using the `‘ctmm’` R package), and “Report” (a summary page of all inputs, outputs, and final recommendations).

The application consists of four main sections, organized as easy-to-navigate tabs seen in the left sidebar. Once the user has set their data source and research question(s), the sidebar will automatically subset to only the required steps. We advise users to follow along the guided tutorial available in the “Home” tab, until they feel confident with the application’s workflow. Each run-through of the entire workflow may take a few minutes up to several hours to run; computation bottlenecks arise from the `ctmm::ctmm.select()` and `ctmm::speed()` functions, particularly if the absolute sample size exceeds 10^3^ locations, or for very short sampling intervals.

### 3.1. Importing movement data

There are three data source options available in ‘`movedesign`’: (1) Import data, (2) Select data, and (3) Simulate data. For the first option, and prior to using ‘`movedesign`’, users should find an animal tracking dataset for their intended study species (*e*.*g*., from Movebank, published studies, requesting directly from data owners), preferably in a similar environment to the focal study site (*e*.*g*., natural *versus* human-dominated environment); if this is not available, search for species with a similar physiological and behavioral phenotype (*e*.*g*., the same genus or family, occupying the same ecological niche, similar body size). As the outputs from ‘`movedesign`’ depend heavily on the extracted species parameters from pre-existing datasets, this step is crucial to ensure the interpretability and reliance of all subsequent steps. Correctly importing data into the ‘`movedesign`’ application requires that datasets conform to Movebank naming conventions (http://www.Movebank.org/). We recommend that users first upload their datasets (privately if needed) to Movebank, as this will facilitate data preparation. For the second option, the user can select one of the seven example species available within the ‘`ctmm`’ package and extract their parameters, which serve as worked examples, but may also help inform animal tracking studies of species with similar movement behaviors to those listed. For the third option, the user can simulate an animal tracking dataset from scratch by inputting model parameters (*τ*_*p*_, *τ*_*v*_) and associated measures (*σ*_*p*_, *σ*_*v*_) directly; if chosen, this option requires previous knowledge of the study species’ movement behavior, and an educated guess regarding their species parameters. For range-resident species, *τ*_*p*_ is generally on the order of days to weeks, while *τ*_*v*_ is generally on the order of minutes to hours (Noonan, Tucker, et al., 2019); however, these parameters are likely to differ between species, populations, or individuals.

For options one and two (“Import data” and “Select data”), the application checks if datasets are valid: specifically, if there is still a signature of the animal’s position or velocity autocorrelation parameter. For example, if the user is interested in speed estimation but provides only a coarsely-sampled tracking dataset (Δ*t* ≫ *τ*_*v*_) then the property of continuous velocities is not supported by the data and the resulting movement model is fractal (*i*.*e*., we cannot proceed with speed estimation). Data visualization boxes are also available to assist in additional validation steps that cannot be done automatically, such as variograms to confirm range residency if the goal is home range estimation (Fleming et al., 2014a), or plots to identify outliers (which should be removed and the data re-uploaded before proceeding).

For option three (“Simulate data”), the application will simulate autocorrelated tracking data using an isotropic Ornstein-Uhlenbeck with foraging (OUF) Gaussian process, which incorporates both correlated velocities and constrained space use. This OUF model is also the most frequently selected across modern GPS tracking datasets (Noonan, Tucker, et al., 2019).

### 3.2. Choosing sampling design

The Device section allows for the simulation of a new dataset, conditioned upon previously extracted parameters, and based on the sampling design of the user’s choice. First, users are prompted to input their desired device type (GPS/Satellite loggers or VHF transmitter) as appropriate sampling design is dependent on the device’s battery life: for GPS/Satellite loggers, battery life depends on both duration and frequency, while for VHF transmitters, battery life depends only on duration. For the GPS option, if users do not intend to visualize the trade-off curve and already have a specific sampling design in mind, they can uncheck the ‘Select from plot’ to ‘Set regime manually’ located in the footer of the Device settings box (this is equivalent to setting VHF transmitter as the device of choice).

The application will once again validate all current inputs, and inform the user of the expected run time of the simulation step. For home range estimation, we recommend that the user check the range residency assumption through the variogram in the data visualization boxes. The user can also check the effective sample sizes extracted from the new model fit, which will assist in decision making (to continue with the current sampling design, or to make further adjustments before proceeding).

### 3.3. Running estimators and building the report

In the Analyses section, there will be one or two tabs available to the user, based on the chosen outputs: (1) Home range, and/or (2) Speed & distance. Each tab will run an estimator on the simulated dataset, created during the previous section. For each output, there will be two sets of values in the main output box: “Estimate”, which is the point estimate followed by the 95% confidence intervals (CIs), and “Expected error”, which is the relative error (in %) of the point estimate (and of the 95% CIs) in relation to the expected values. The user can utilize these outputs (as well as effective sample sizes) to plan an appropriate sampling duration and interval so the confidence intervals are sufficiently narrow and relative errors acceptably low: typically, a more reliable home range estimate requires a longer sampling duration than the one specified, while a more reliable speed and distance estimation requires shorter sampling intervals. However, we advise caution in two scenarios for speed and distance estimation: (1) producing a CTSD estimate at Δ*t* > 3*τ*_*v*_ does not guarantee a meaningful output, and (2) Δ*t* ≪ *τ*_*v*_ may only yield a marginal benefit over Δ*t* < *τ*_*v*_ while markedly increasing computation time (see Noonan, Fleming, et al., 2019 for more details). Users can then navigate to the last section, the “Report” tab, to see a summary of all previous inputs and outputs, and the final recommendations regarding home range and/or speed and distance estimation.

## 4. Case study

To present the application’s key features, we used the African Buffalo (*Syncerus caffer*) data available within the ‘`ctmm`’ `R` package as a case study. These individuals were tracked in Kruger National Park, South Africa (Cross et al., 2009). Our objectives were to reliably estimate home range area and speed/distance traveled (Figure 2) using GPS loggers for a hypothetical tracking project with the same species in a similar study site. We choose a GPS model with a maximum lifespan of two years if four new locations were collected every day (Δ*t* = 1 fix every 6 hours; Figure 3).

**Figure 2.**
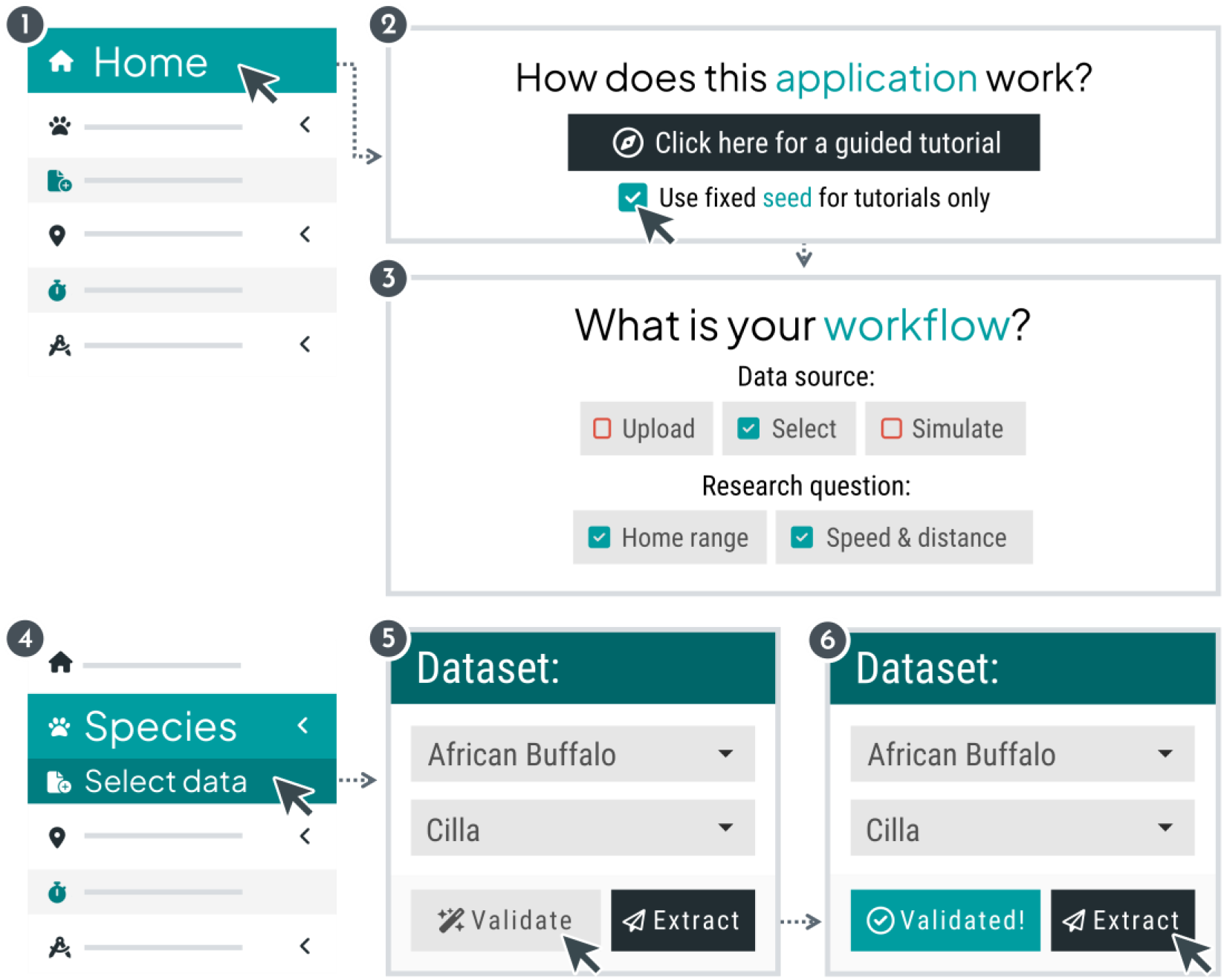
Initial steps in the `‘movedesign’` Shiny application workflow (“Home” and “Species” tabs). For the case study presented here, set a fixed seed to reproduce all outputs, the correct data source, both research questions, and the stated species and individual from the list (before validation and extraction).

**Figure 3.**
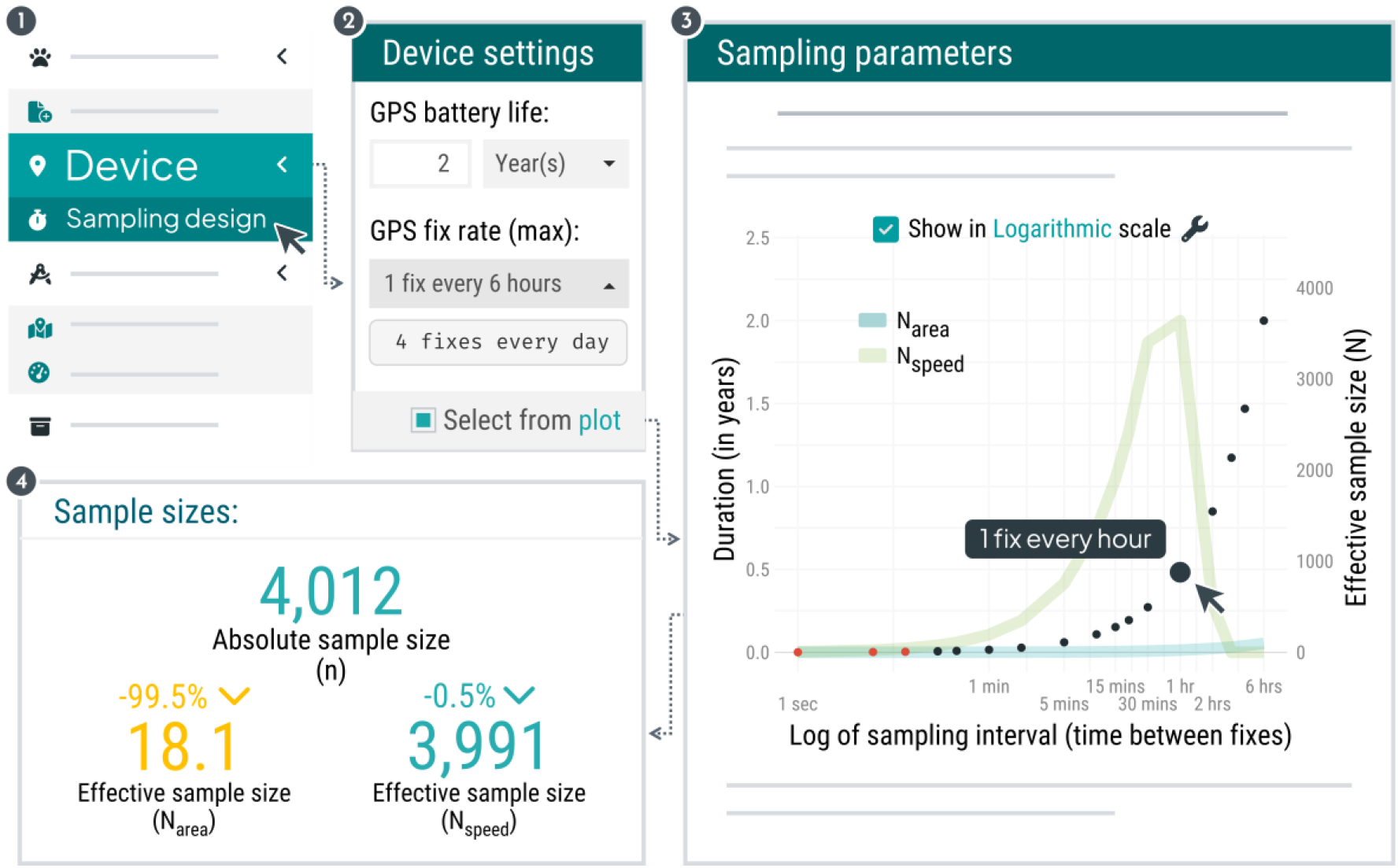
Example outputs shown in the “Device” tab, including the simulated trade-off between sampling duration and sampling interval (shown in logarithmic scale) for the specified device settings, with the chosen interval selected from the plot (1 fix every hour), and the absolute and effective sample sizes extracted from the new model fit.

We selected the individual “Cilla” for parameter extraction; once validated, we extracted parameters from the fitted OUF anisotropic model for a *τ*_*p*_ of approximately 7.5 days (CI: 4.4 – 12.7), and a *τ*_*v*_ of 42 minutes (39.6 – 44.7). We plotted our sampling design options based on the chosen tracking device parameters (Figure 3), which revealed that a rough estimate of *N*_*area*_ for any sampling interval was substantially smaller the *N*_*speed*_ (as expected, since *τ*_*p*_ ≫ *τ*_*v*_). As our focus was both home range and speed/distance estimation, we wanted to maximize *N*_*area*_ and *N*_*speed*_, so we selected a sampling interval of one hour (which sets our sampling duration to 5.7 months, due to the battery/resolution trade-off discussed earlier). Once we successfully validated and ran our new simulation, our sample sizes were *n* = 4,012, *N*_*area*_ = 18.1 and *N*_*speed*_ = 3,991.

With an appropriate movement model available, the next step was to estimate home range area, mean speed and total distance traveled (Figure 4). Once the final report was built, the AKDE estimates showed high uncertainty, while the CTSD estimates did not. Based on these results and the effective sample sizes, we determined that our sampling design was adequate for speed/distance estimation, but could be insufficient for home range (Figure 5). After exploring further sampling designs for our choice of GPS model, we decided upon one fix every 2 hours for a sampling duration of 11 months, reducing the uncertainty associated with home range area ―although at the cost of increased uncertainty in speed/distance estimation (follow along the guided tutorial in the “Home” tab for this particular workflow).

**Figure 4.**
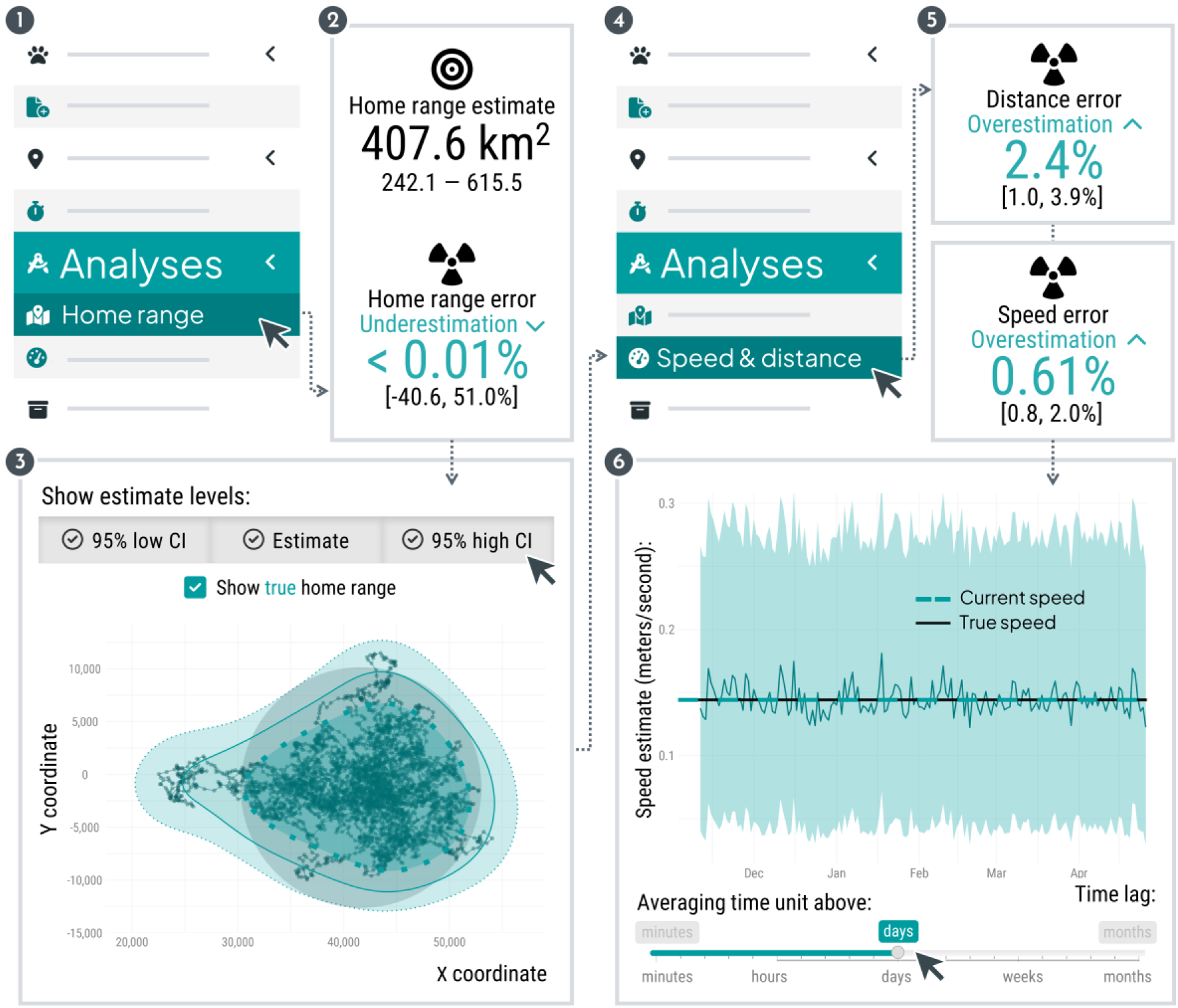
Example outputs shown in the “Analyses” tabs, including the point estimates (and corresponding confidence intervals below the point estimate in small, black text) and/or expected errors associated with both home range and speed/distance. We can also visualize the home range estimate (with the true 95% area for comparison), and the instantaneous speed estimates at different time scales —other visualizations (such as those related to distance) are not shown here but are available within the app.

**Figure 5.**
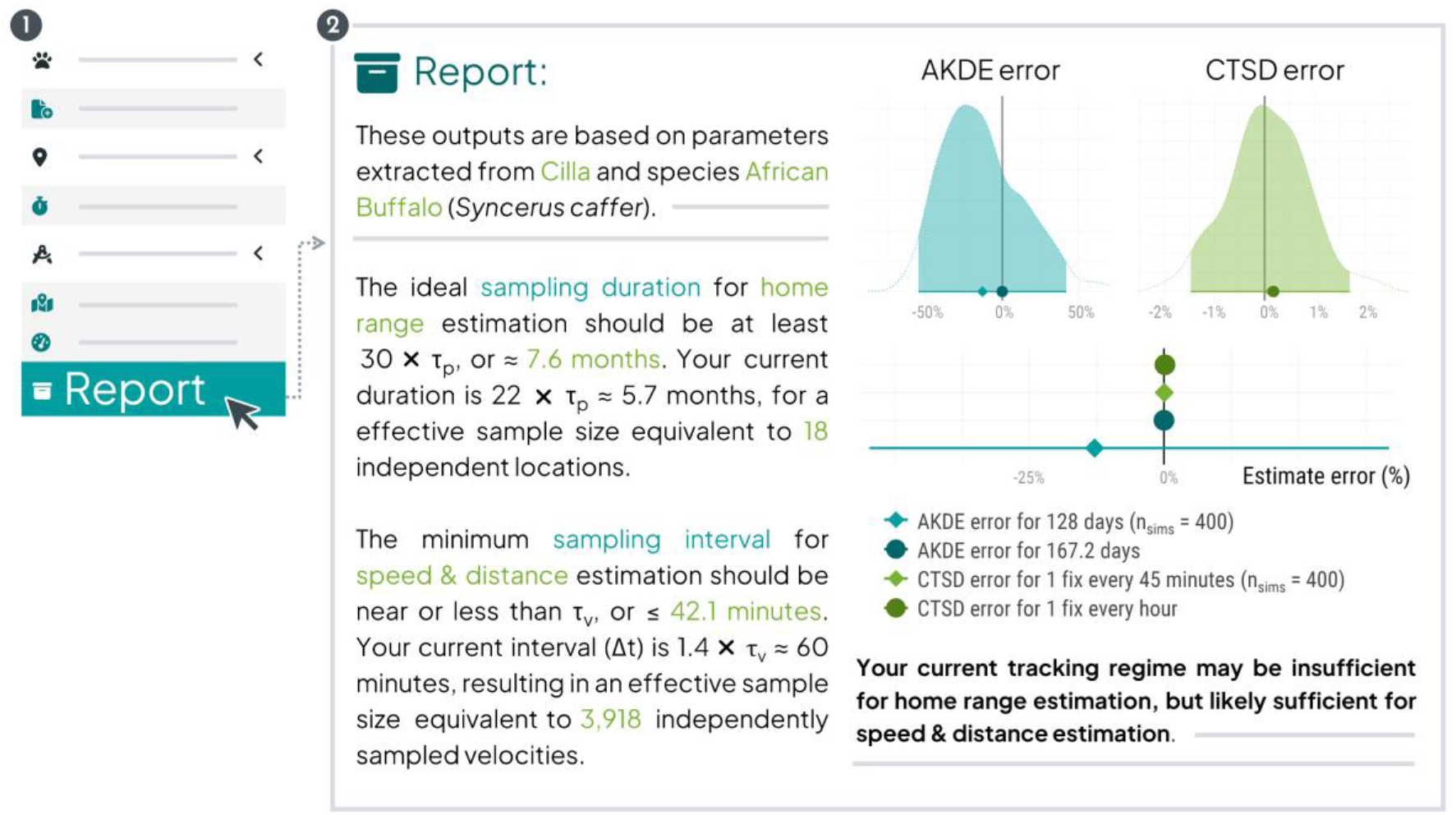
Final report for a workflow that requested both home range and speed/distance as the research questions, for a hypothetical tracking study of the African buffalo (*Syncerus caffer*) species with a sampling duration of 5.7 months and a sampling interval of one hour —based on a GPS model with a maximum battery life of 2 years if the sampling interval was set to 6 hours.

## Final considerations

Data collection and sampling design optimization allows researchers to better address key ecological questions, as conclusions drawn from lower-resolution data may not be appropriate to detect fine-scale movement properties, while higher-resolution datasets may still be unsuitable to detect patterns at larger spatial or temporal scales (*e*.*g*., space-use), if the sampling duration is too short (Nathan et al., 2022). Our workflow allows users to evaluate a wide range of potential sampling designs, which can then serve as a solid foundation for future tracking projects, or even the evaluation of on-going and published studies. One limitation of our study is that our GPS simulations do not consider the case of solar tags. As a workaround, users can test the minimum expected battery life or the duration of the entire tracking study. Our application also assumes, for the purposes of home range estimation, that the intended study species is range-resident; if users plan to track migratory species, they should consider assessing resident periods (before and/or after migration, if applicable) as separate study design scenarios, each requiring their own assessment. We recommend that users employ the same methods during the study planning phase (facilitated here through the ‘`movedesign`’ application) and the final analyses after data collection to ensure similar effective sample sizes. We are continuously working to improve the application’s code, and have plans to explore new use cases and address more challenging sampling design questions.

## Data Availability statement

The empirical dataset used in the manuscript is openly accessible: the African buffalo tracking data are archived in the MoveBank Data Repository (Cross et al., 2016) and partially included in the ctmm package (Fleming & Calabrese, 2022). Source code of the ‘`movedesign`’ application is available on GitHub (https://github.com/CASUS/movedesign), and release version 0.1.3 has been archived on Zenodo (Silva, 2023).

## Acknowledgements

This work was partially funded by the Center of Advanced Systems Understanding (CASUS), which is financed by Germany’s Federal Ministry of Education and Research (BMBF) and by the Saxon Ministry for Science, Culture and Tourism (SMWK) with tax funds on the basis of the budget approved by the Saxon State Parliament. C.H.F. and J.M.C. were supported by NSF IIBR 1915347.

## Conflict of interests

The authors declare no conflicts of interest.

## Authors’ contributions

I.S., C.H.F., M.J.N., and J.M.C. conceived the ideas; I.S. designed and developed the application; J.M.C. and I.S. led the drafting the manuscript; I.S., C.H.F., M.J.N., W.F.F., and J.M.C. edited the draft. All authors reviewed the application, contributed critically to the manuscript and gave final approval for publication.

## Notes

https://github.com/CASUS/movedesign

### Competing Interest Statement

The authors have declared no competing interest.

### Summary of Updates

Manuscript and figures updated.

