## Appendix S1 for "movedesign: Shiny R app to evaluate sampling design for animal movement studies"

To visualize GPS battery life trade-off (sampling duration *versus* sampling interval), we simulated GPS battery life as a three-parameter log-logistic function with lower limit 0;

$$b = \frac{b_{max}}{1 + e^{p(\log(\frac{1}{\Delta t}) + \log b_{50})}} \quad [1]$$

where $1/\Delta t$ is the fix rate (1/sampling interval), $b$ is the predicted battery life, $b_{max}$ is the maximum battery life, and $p$ governs the steepness of the decay in battery life at $b_{50}$ (50% of the battery life capacity).

This function is based on the lifespan of several GPS logger models provided by two manufacturers of animal tracking devices: Lotek (Lotek.com) and Advanced Telemetry Systems (ATStrack.com). The exact decay rates and parameters are likely to vary between companies and models, but the present configuration only requires users to provide two parameters: the maximum expected battery life (how long the GPS is expected to last), and the maximum GPS fix rate (the sampling interval between new recorded fixes available at the duration above). Users can then match a trade-off curve to the expected values for a specific model reported by the manufacturer of their choice. The underlying pattern is that sampling duration decreases as the user sets a higher fix rate (shorter sampling interval), but the decay rate slows down for very short sampling intervals. Specifically, if the lag between fixes is very short, the battery power needed to determine the next location decreases (Moriarty & Epps 2015; McGregor *et al.* 2016).
